# Attenuating LRRK2 activity ameliorates progerin-induced aging phenotypes in HGPS models and during physiological aging

**DOI:** 10.64898/2026.08.04.742751

**Authors:** Lukas Mann, Rosa Herrera Rodriguez, Felix van der Walt, Kyle R Brimacombe, Sheraz Sadouki, Tom Misteli, Nard Kubben, Jan Padeken

## Abstract

Hutchinson-Gilford progeria syndrome (HGPS) is an ultra-rare premature aging disorder caused by progerin, a truncated lamin A variant generated by a silent *de novo* mutation activating a cryptic splice site in *LMNA*. The resulting morphological, epigenetic, genomic, and proteostasic defects closely recapitulate some hallmarks of cellular aging. Here, we identify the Parkinson’s disease-associated kinase LRRK2 as a critical regulator of HGPS pathology and physiological aging. Rab29-mediated LRRK2 hyperactivation exacerbates progerin-induced cellular aging, whereas *LRRK2* knockdown or overexpression of its opposing phosphatase, PPM1H, ameliorates progerin-induced defects. Progerin-expressing cells exhibit altered intracellular trafficking, which is regulated by LRRK2 and links diverse aging hallmarks. Consistent with these findings, reducing LRRK2 levels mitigates cellular aging phenotypes in physiologically aged cells, and loss of the *C. elegans* ortholog *lrk-1* preserves aging-associated loss of motility and extends organismal lifespan. Together, our findings establish LRRK2 as a central node in cellular aging and position it as a potential therapeutic target for aging-related defects in both HGPS and physiological aging.

## Introduction

Aging is the predominant risk factor underlying the majority of deaths caused by cardiovascular diseases, cancer, and neurodegenerative disorders ^1^. To develop efficient therapeutic strategies against common ageing-associated diseases (AADs), we need to understand the mechanisms that drive aging on a systemic and molecular level. Premature aging diseases such as Hutchinson-Gilford progeria syndrome (HGPS) have proven to be powerful models for studying defects associated with physiological aging and AADs.

HGPS is an ultra-rare genetic disorder that is caused by a silent *de novo* mutation in the *LMNA* gene, which activates an intronic cryptic splice site, resulting in the expression of a truncated lamin A referred to as progerin ^2,3^. Both progerin and wild-type (WT) lamin A are farnesylated and undergo initial proteolytic cleavage steps. However, progerin lacks 50 amino acids at the C-terminus, including the endoproteolytic cleavage site required for a terminal cleavage step mediated by ZMPSTE24, an ER–associated metallopeptidase required for prelamin A maturation ^2–4^. Unlike wild-type lamin A, progerin retains its farnesylated C-terminal CAAX-box motif, which permanently tethers progerin to the nuclear membrane, where it accumulates, interfering with normal nuclear lamina function ^3,5^. HGPS cells exhibit drastic alterations in nuclear morphology, accumulation of DNA damage, loss of heterochromatin, telomere shortening, but also non-nuclear metabolic and proteostatic defects, oxidative stress, and eventual premature senescence ^6^. HGPS thereby resembles a multitude of defects also observed during physiological aging ^6^. Interestingly, low levels of progerin have been detected in various cell types during physiological aging, suggesting that progerin may contribute to cellular aging beyond HGPS ^7,8^. Although the ability of progerin to induce cellular aging in HGPS is well established, many of the underlying mechanisms and, importantly, points of intervention remain poorly understood.

Despite the missing organismal context that is required to fully capture all aspects of aging, cell-based models have provided critical insights into HGPS pathobiology. Especially, an inducible cell-based system in which progerin expression recapitulates several premature aging phenotypes within days has provided a robust screening platform to identify pathways that drive progerin-induced cellular aging ^9^. Evidence from both transgenic cell-based model systems and primary HGPS fibroblasts indicates that complex proteotoxic alterations are a driver of HGPS pathology ^10–12^. For example, progerin forms aggregates at the nuclear periphery and sequesters NRF2, a master regulator of the cellular anti-oxidative response, thereby impairing its natural cellular function ^11^. Progerin also inhibits UBC9 nuclear localization, causing the collapse of the Ran gradient and disrupting the nuclear import of large protein complexes ^13,14^. In addition, inner nuclear membrane (INM) proteins like emerin and SUN1 are aberrantly retained with progerin in the ER after mitosis in HGPS cells ^15^. These findings indicate that the effects of progerin are not limited to the nucleus. Indeed, progerin-expressing cells exhibit compromised proteasomal and autophagic flux, lysosomal dysfunction, ER stress, alterations in ER and Golgi morphology, and progeroid mouse models reveal defects in protein secretion and secretory trafficking ^16–21^. Notably, many of the proteotoxic defects in HGPS are also observed in physiological aging and AADs, such as the neurodegenerative Parkinson’s Disease (PD) ^22^. Importantly, endosomal trafficking is broadly impaired in PD, with patients with familial PD carrying mutations in LRRK2, VPS35, SNCA, DNAJC6, SYNJ1, DNAJC13, GBA, ATP13A2, and VPS13C ^23,24^.

One of the prominent genes associated with familial PD is LRRK2 ^25^. Mutations that enhance the activity of the S/T kinase LRRK2 are the most common cause of familial PD ^25^. LRRK2 is a key regulator of the endolysosomal trafficking pathway, acting primarily through phosphorylation of a subset of Rab GTPases and thereby modulating their activity and interaction with downstream effectors ^26,27^. Cells expressing hyperactive LRRK2 mutants exhibit widespread proteotoxic collapse, including protein aggregation, impaired autophagy and lysosomal function, impaired synaptic vesicle endocytosis, and fragmentation of the Golgi apparatus ^28–32^. Therefore, in the context of PD, LRRK2 has emerged as a promising therapeutic target, with clinical trials evaluating the beneficial effects of LRRK2 kinase inhibitors ^33^.

Here we identify LRRK2 as an important regulator of HGPS pathology and physiological aging. Reduction of LRRK2 levels mitigates the cellular aging phenotypes in progerin-expressing fibroblasts and HGPS patient cells. In contrast, overexpression of the LRRK2 activator Rab29 and knockdown of the phosphatase PPM1H, which counteracts LRRK2 activity, result in accelerated cellular aging phenotypes. Our work identifies LRRK2 as a potential therapeutic target for both physiological and progerin-driven cellular aging, while establishing impaired membrane trafficking as a fundamental mechanism underlying progerin-induced cellular pathology.

## Results

### *LRRK2* knockdown ameliorates progerin-induced cellular aging

Kinases are a highly validated drug target class, and they are established modulators of aging, exemplified by MAPK and GSK-3 signaling ^34,35^. We thus used a kinome-wide screen to identify potentially druggable regulators of progeria-induced premature aging. The RNAi screen targeted 704 human genes using three independent siRNAs per construct to identify kinases that drive cellular aging pathologies in a well-established human dermal fibroblast cell line that allows for the rapid induction of GFP–progerin upon addition of doxycycline ^9,11^. The system provides tight control of GFP-progerin expression and induces several quantifiable cellular aging phenotypes 96 hours post-induction, including reduction of lamin B1 levels and increased DNA damage ^9^. Therefore, we assessed these three independent cellular aging parameters by high-throughput microscopy coupled to automated image analysis: GFP-progerin levels, loss of lamin B1 (anti-lamin B1), and accumulation of DNA damage foci (anti-γH2AX; Fig. 1A, Supplementary Table 1). Consistent with prior reports, knockdown of *MAPK14* (p38α) rescued lamin B1 levels (p **=** 0.0168), but modestly increased γH2AX foci (p = 0.00902), without altering GFP–progerin expression (p = 0.152) (Fig. S1A) ^36^. Similarly, *MAPK11* (p38β) knockdown rescued lamin B1 loss in progerin-expressing cells (p = 0.05) but exhibited heterogeneous siRNA-dependent responses, with no consistent effects on GFP–progerin levels or γH2AX accumulation (Fig. S1A) ^37^. To leverage the screen despite variability in the siRNAs, we evaluated the effect of individual siRNAs independently. In the screen, a total of 20 hits were found that affected all three readouts, and 9 hits rescued loss of lamin B and γH2AX accumulation. Intriguingly, a top 5 hit was the PD-related LRRK2 kinase, reducing each of the three measured cellular aging parameters (Fig. 1B, S1B).

**Figure 1:**
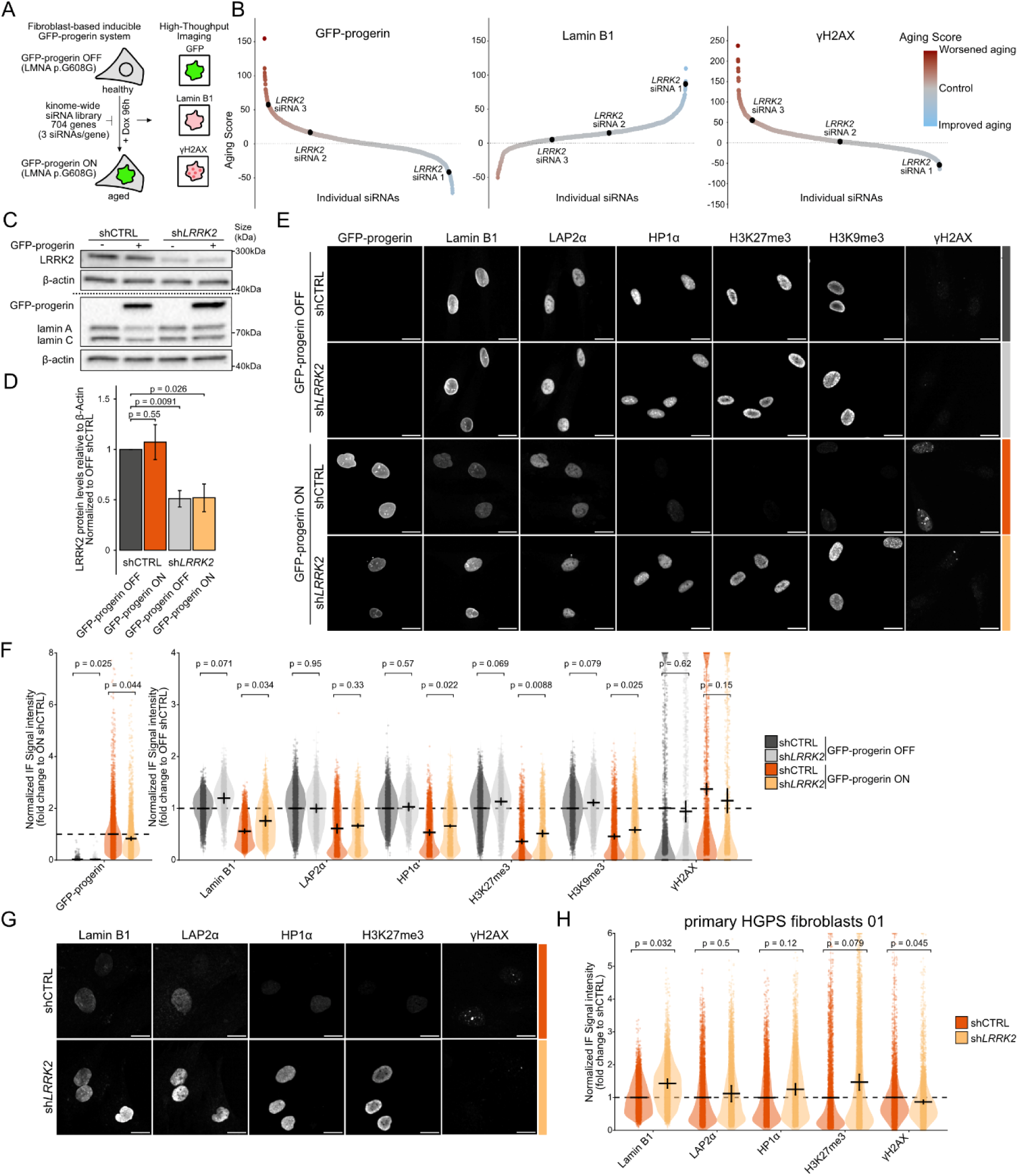
Reduction of LRRK2 alleviates aging-related cellular phenotypes in the inducible HGPS model and primary HGPS patient cells. A: Schematic of the RNAi screen in a human dermal fibroblast-based system with inducible GFP–progerin expression. B: Waterfall plots depicting RNAi screen results for the evaluated aging parameter. Each dot represents an individual siRNA, aging score indicates the change relative to control siRNA-treated cells. siRNAs are ranked by aging score. siRNAs targeting *LRRK2* are highlighted. C, D: Representative Western blot (C) of LRRK2 and lamin A/C with β-actin as a loading control, and quantification (D) of LRRK2 levels normalized to β-actin in progerin-inducible fibroblasts expressing control or *LRRK2*-targeting shRNA. GFP-progerin-inducible fibroblasts were incubated with 250 ng/ml doxycycline for 96 h. Values represent means ± SD (N=3). Statistical significance was assessed using a paired two-tailed Student’s t-test. E, F: Representative immunofluorescence images (E) and quantification (F) of cellular aging markers in progerin-inducible fibroblasts expressing control or *LRRK2*-targeting shRNA. GFP-progerin-inducible fibroblasts were incubated with 250 ng/ml doxycycline for 96 hours. Scale bar: 20 μm. Values represent means ± SD (N=3). Statistical significance was assessed using a paired two-tailed Student’s t-test. G, H: Representative immunofluorescence images (G) and quantification (H) of cellular aging markers in HGPS patient fibroblasts expressing control or *LRRK2*-targeting shRNA. Scale bar: 20 μm. Values represent means ± SD (N=3). Statistical significance was assessed using a paired two-tailed Student’s t-test.

To orthogonally validate the potential effects of *LRRK2* siRNAs, we performed shRNA-mediated *LRRK2* knockdown in two independent progerin-inducible fibroblast cell lines with distinct doxycycline-responsive promoters and evaluated a more comprehensive panel of cellular aging markers. The shRNA-mediated knockdown reduced LRRK2 protein levels by ∼50% (Fig. 1C and D). GFP-progerin induction itself did not change either LRRK2 RNA or protein levels, suggesting that progerin does not drive cellular aging through transcriptional or translational upregulation of LRRK2 (Fig. 1C and D, S2A). In both cell lines, *LRRK2* knockdown reduced progerin levels and partially rescued progerin-associated loss of nuclear envelope and lamina proteins lamin B1 and LAP2α, decrease of heterochromatin markers HP1α, epigenetic marks H3K27me3, and H3K9me3, and accumulation of DNA damage indicated by γH2AX foci formation in progerin-expressing cells (Fig. 1E and F, S2B) ^9^. In the absence of progerin, loss of LRRK2 had no significant effects on these markers, except for a mild increase in lamin B1 (Fig. 1E and F, S2B). To test if a reduction of LRRK2 levels also has beneficial effects in HGPS patient cells, we knocked down *LRRK2* in three independently derived primary HGPS fibroblast lines using two newly designed shRNAs in addition to the original shRNA 01 (subsequently referred to as sh*LRRK2*) to rule out potential off-target effects (Fig. S2C). Knockdown of *LRRK2* using both shRNAs individually consistently improved cellular aging across the 3 primary HGPS fibroblast lines, indicated by increased levels of lamin B1 and LAP2α, alongside recovery of HP1α and H3K27me3 levels, and a reduction of γH2AX foci (Fig. 1G and H, S2D and S2E). This validates LRRK2 as a candidate from the screen that ameliorates cellular aging in progerin-expressing cells. These results also show that reducing LRRK2 levels benefits not only the acute onset of progerin pathology but also cells with an already established cellular aging phenotype.

### Overexpression of the LRRK2-opposing phosphatase PPM1H ameliorates progerin-induced cellular aging

LRRK2 is a key regulator of endolysosomal trafficking, acting primarily through phosphorylation of a subset of Rab GTPases, which function as molecular switches regulating the budding, transport, tethering, and fusion of vesicles ^26^. As reducing LRRK2 levels counteracted the detrimental effects of progerin, we hypothesized that increasing the LRRK2 activity could drive cellular aging on its own or in combination with progerin. While simply overexpressing LRRK2 did not result in a measurable effect on the cellular aging markers tested (Fig. S3A-D), a mild cellular aging phenotype was induced when we specifically increased LRRK2 activity by overexpression of Rab29 (Fig. 2A and B), another PD-linked protein that functions as a well-characterized activator of LRRK2 and mediates both LRRK2 kinase activation and its recruitment to the Golgi apparatus ^38,39^. Overexpression of Rab29 in progerin-expressing cells accelerated the progerin-induced phenotypes of most markers, except for a modest reduction in γH2AX foci without increasing GFP-progerin levels (Fig. 2A and B).

**Figure 2:**
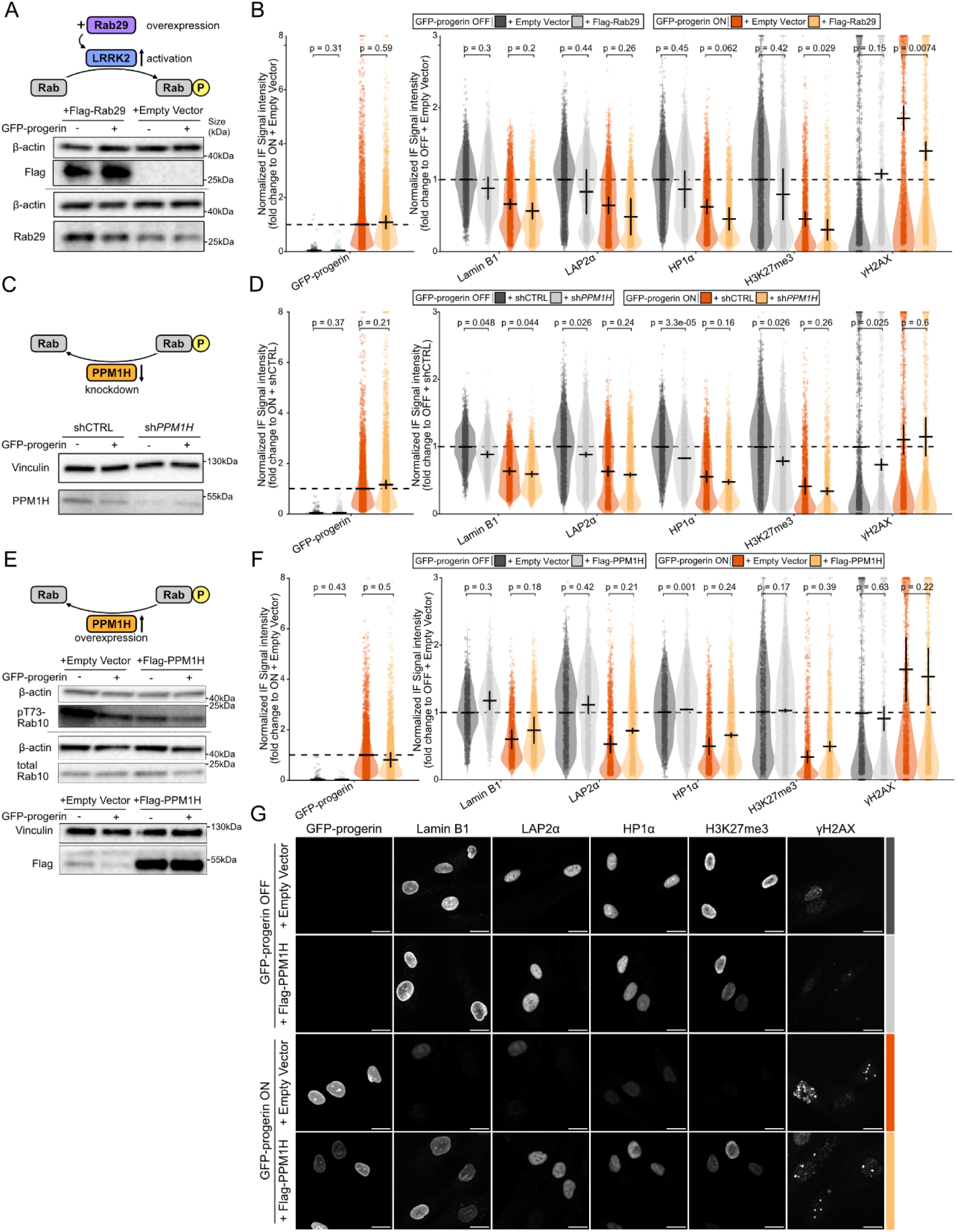
Progerin-induced cellular aging is mediated by the LRRK2/PPM1H pathway. A: Representative Western Blot of Flag-tag and Rab29 with β-actin as a loading control in progerin-inducible fibroblasts expressing an empty vector or Flag-Rab29. B: Immunofluorescence quantification of cellular aging markers in progerin-inducible fibroblasts expressing an empty vector or Flag-Rab29. Values represent means ± SD (N=3). Statistical significance was assessed using a paired two-tailed Student’s t-test. C: Representative Western Blot of PPM1H with β-actin as loading control in progerin-inducible fibroblasts expressing control or *PPM1H*-targeting shRNA. D: Immunofluorescence quantification of cellular aging markers in progerin-inducible fibroblasts expressing control or *PPM1H*-targeting shRNA. Values represent means ± SD (N=3). Statistical significance was assessed using a paired two-tailed Student’s t-test. E: Representative Western Blot of Flag-tag and phosphorylated Rab10 (pT73 Rab10) relative to total Rab10 with β-actin as loading control in progerin-inducible fibroblasts expressing an empty vector control or Flag-PPM1H. F, G: Representative Immunofluorescence images (F) and quantification (G) of cellular aging markers in progerin-inducible fibroblasts expressing an empty vector or Flag-PPM1H. Scale bar: 20 μm. Values represent means ± SD (N=2). Statistical significance was assessed using a paired two-tailed Student’s t-test.

If LRRK2 kinase activity is driving the premature aging phenotype, loss of PPM1H, the phosphatase known to counteract LRRK2 phosphorylation, should mimic the effect of Rab29 overexpression ^40^. Indeed, knockdown by shRNA of *PPM1H* induced a cellular aging phenotype in the absence of GFP-progerin, except for a modest reduction in γH2AX foci (Fig. 2C and D, S3E and F). Interestingly, *PPM1H* knockdown did not further accelerate GFP-progerin-induced cellular aging, suggesting that progerin may drive cellular aging through the LRRK2/PPM1H pathway. Conversely, we hypothesized that overexpression of PPM1H would phenocopy the effects of *LRRK2* knockdown and rescue progerin-induced cellular aging defects. To confirm activity of the overexpressed FLAG-PPM1H, we evaluated levels of phosphorylated Rab10 (pT73-Rab10), a canonical substrate of LRRK2 and PPM1H ^40^. As expected, phospho-Rab10 levels significantly decreased following PPM1H overexpression (Fig. 2E, S3G). PPM1H overexpression indeed rescued progerin-induced senescence and partially restored nuclear LAP2α and lamin B1 in uninduced cells (Fig. 2F, G). Together, these observations argue that the beneficial effects of *LRRK2* knockdown are mediated through its canonical role in endolysosomal trafficking and suggest that LRRK2 activity, rather than abundance, is responsible for the observed cellular aging phenotypes.

To test if the known substrates of LRRK2 contribute to the effect on cellular phenotypes, we overexpressed WT, constitutively active (CA), and dominant-negative (DN) mutants of Rab10, Rab12, Rab35, and Rab43. These are canonical substrates of LRRK2, alongside Rab9A as a non-canonical control ^41^. Together, these Rab GTPases are important regulators of ER-to-Golgi trafficking and secretion, endocytosis, recycling pathways, and lysosomal function (Fig. S3H) ^41^. In the absence of GFP-progerin, overexpression of DN Rab10 or CA Rab35 induced cellular aging compared to cells expressing an empty control vector (Fig. S3I and J). Only CA Rab35 accelerated GFP-progerin-induced cellular aging. While dysregulation of Rab proteins has multiple pleiotropic effects, precluding the identification of a clear driver pathway that mediates the cellular aging effects of LRRK2, these results further suggest an important role of endolysosomal trafficking in cellular aging and indicate that LRRK2 likely drives cellular aging through its canonical function in regulating a subset of Rab GTPases ^26^.

### Progerin induces defects in secretory, nucleocytoplasmic, and endolysosomal trafficking

Because modulation of the LRRK2–PPM1H axis and its substrates directly influenced progerin-driven cellular aging, we asked if, and how, protein trafficking is affected in progerin-expressing cells. To test whether progerin expression causes widespread structural damage in the protein trafficking pathway, we quantified the organization of the Golgi apparatus/complex and the ER. To quantify overall ER morphology, we stained for the ER-resident chaperone BIP but found no visible change upon progerin induction (Fig. S4A and B). Hyperactive LRRK2 can, under specific circumstances, trigger Golgi fragmentation ^38,42,43^. Therefore, we stained for GRASP65, a membrane protein specific to the Golgi apparatus, to evaluate if Golgi morphology is affected in progerin-expressing cells. Quantification revealed a non-significant 10% decrease in Golgi apparatus volume, accompanied by a very minor ∼10% increase in fragmentation in progerin-expressing cells (Fig. S4A and B). To explore whether secretory pathway dysfunction could in principle contribute to age-associated nuclear defects, we treated cells with GolgiStop, a transport inhibitor containing the ionophore monensin that disrupts the pH gradient in the Golgi apparatus and thereby prevents movement of proteins from cis to trans Golgi apparatus and impairs endolysosomal function ^44^. Pharmacological disruption of Golgi function induced a reduction in nuclear roundness compared to the pronounced nuclear deformation observed in progerin-expressing cells, suggesting that impaired secretory trafficking can compromise nuclear morphology (p-value: 0.032; Fig. S4C and D). Quantification of an extended cellular aging marker panel revealed that progerin-expressing cells and cells treated with the GolgiStop inhibitor exhibit a similar cellular aging phenotype, especially loss of heterochromatin and the collapse of the Ran gradient. Notably, the DNA damage response (particularly γH2AX foci) was less pronounced following Golgi apparatus disruption compared to progerin induction (Fig. S4E). These results suggest a connection between Golgi defects and progerin-induced cellular aging phenotypes.

Because disruption of secretory transport phenocopied key cellular aging features, we asked if protein trafficking defects are part of HGPS pathology. To measure defects in protein trafficking, we established the Retention Using Selective Hooks (RUSH) assay to monitor ER-to-Golgi trafficking in progerin-inducible cells ^45^. The RUSH assay comprises a Hook protein fused to streptavidin and stably anchored to a donor compartment, it captures a reporter protein fused to a streptavidin-binding peptide (SBP) and a fluorescence marker. Biotin addition releases the reporter, which can then be transported to its destination compartment (Fig. 3A). Using two distinct Hook/Reporter combinations (Str-STIM1/SBP-mCherry-GPI, Ii-Str/TNFα-SBP-mCherry), we quantified reporter accumulation in the Golgi apparatus over time by fluorescence microscopy ^45^. In the absence of biotin, the GPI reporter was uniformly distributed throughout the cell, while accumulation in the Golgi apparatus became evident 15 minutes after biotin treatment and plateaued after 60 minutes in uninduced control cells (Fig. 3B and C). Upon GFP-progerin induction, reporter accumulation in the Golgi apparatus was significantly delayed and only reached comparable accumulation after 60 minutes of biotin treatment (Fig. 3B and C). The TNFα reporter exhibited cytosolic localization in the absence of biotin (Fig. 3D). Upon Biotin addition, the reporter rapidly accumulated in the Golgi apparatus and peaked at 15 minutes, declining by 60 minutes (Fig. 3D and E). In GFP-progerin-expressing cells, Golgi apparatus accumulation was similarly delayed compared to the progerin OFF state, continuously increasing at 60 minutes (Fig. 3D and E). Notably, the TNFα reporter colocalized with GFP-progerin and accumulated in the nucleus in progerin-expressing cells, with 6.7% of cells showing a strong nuclear reporter signal even without biotin treatment, suggesting misdirection of cargo from the ER to the nucleus in progerin-expressing cells (Fig. S5A-C).

**Figure 3:**
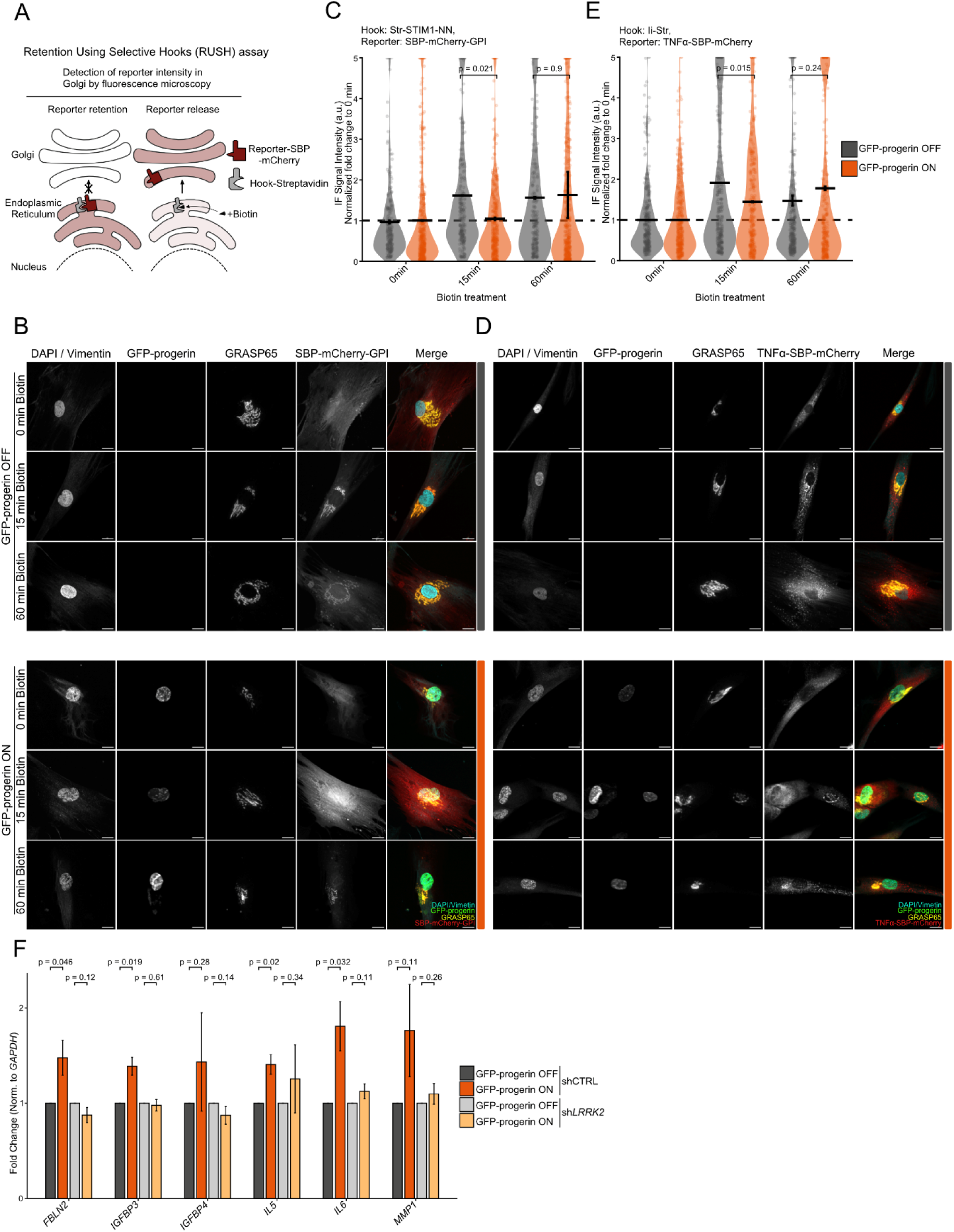
Progerin impairs ER-to-Golgi protein trafficking and a senescence-associated secretory phenotype (SASP), attenuated upon LRRK2 knockdown. A: Schematic of the RUSH assay to monitor protein trafficking from the Endoplasmic Reticulum to the Golgi apparatus. B, C: Representative Immunofluorescence images (B) and quantification (C) of the RUSH reporter in progerin-inducible fibroblasts expressing Str-STIM1-NN as a hook and SBP-mCherry-GPI as a reporter. Scale bar: 20 μm. Values represent means ± SD (N=2). Statistical significance was assessed using a paired two-tailed Student’s t-test. D, E: Representative Immunofluorescence images (D) and quantification (E) of the RUSH reporter in progerin-inducible fibroblasts expressing li-Str as a hook and TNFα-SBP-mCherry as a reporter. Scale bar: 20 μm. Values represent means ± SD (N=2). Statistical significance was assessed using a paired two-tailed Student’s t-test. F: qPCR analysis of mRNA levels of SASP gene expression in progerin-inducible fibroblasts expressing control or *LRRK2*-targeting shRNA. GFP-progerin-inducible fibroblasts were incubated with 250 ng/ml doxycycline for 96 hours. Gene expression was normalized to *GAPDH* and is relative to the GFP-progerin OFF control for each shRNA condition. Values represent means ± SD. (N=3). Statistical significance was assessed using a paired two-tailed Student’s t-test.

Given the defects in protein transport, especially the misdirection of proteins from the ER to the nucleus, we wondered if progerin expression also affects the import of bona fide nuclear proteins into the nucleus. Interestingly, previous reports suggested a potential abnormal retention of inner nuclear membrane (INM) proteins in the ER and cytoplasm during nuclear envelope reformation post-mitosis^15^. Quantification of immunofluorescence for INM proteins LBR, SUN1, emerin, and SUN2 revealed protein-specific alterations rather than a uniform directional shift in localization (Fig. S5D). Progerin induction reduced the nuclear/cytosolic ratios of LBR and emerin but increased the nuclear/cytosolic ratio of SUN2 and left SUN1 largely unaffected (Fig. S5D). *LRRK2* knockdown did not alter SUN2 distribution but decreased the emerin nuclear/cytosolic ratio, while increasing the ratio for SUN1 (Fig. S5D). Notably, LRRK2 depletion also elevated both nuclear LBR levels and its nuclear/cytosolic ratio (Fig. S5D). To determine whether these localization shifts were due to transcriptional changes, we performed RT-qPCR for *LBR*, *SUN1*, *EMD*, and *SUN2*, which showed no significant alterations in mRNA levels under any of the conditions (Fig S5E). Together, these protein-specific changes in protein localization indicate that neither progerin expression nor LRRK2 loss causes a global defect in inner nuclear membrane protein trafficking or nuclear import but affect individual INM proteins differentially.

While defective transport of INM proteins may directly affect the pathological nuclear events triggered by progerin, LRRK2 also regulates endocytosis ^26^. Interestingly, impaired endocytosis is part of the complex Parkinson’s disease phenotype in dopaminergic neurons expressing hyperactive LRRK2 mutants ^46^. The breakdown of endocytosis is linked to the senescence-associated secretory phenotype (SASP) profile observed in HGPS patients and a driver of senescence in physiological aging models ^47,48^. We confirmed induction of SASP genes by progerin expression in our system (Fig. 3F). *LRRK2* knockdown attenuated the upregulation of most tested SASP factors upon progerin induction (Fig. 3F). Given that endocytic dysfunction is a well-established trigger for senescence, we next tested whether progerin induces defects in endocytosis. Cells were incubated with Dil-LDL (DiI-labeled low-density lipoprotein) as a standard assay to trace endocytic uptake and subsequent transport to late endosomes and lysosomes by co-staining with markers for early endosomes (EEA1), late endosomes (Rab7), and lysosomes (LAMP2) (Fig. S6A). At 30 minutes in uninduced cells, Dil-LDL was primarily localized at early endosomes and late endosomes with reduced localization at lysosomes. Upon GFP-progerin induction, Dil-LDL signal intensity was reduced across all cellular compartments (Fig. S6B and C). At 2 hours, in uninduced cells, Dil-LDL localized to both late endosomes and lysosomes. In contrast, progerin-expressing cells exhibited reduced signal across compartments for Dil-LDL at 2 hours, with the highest localization in late endosomes. These findings suggest impaired Dil-LDL endocytic uptake upon progerin expression as well as delayed maturation from late endosomes to the lysosome (Fig. S6B and C). Taken together, we observe that progerin expression induces widespread defects in cellular trafficking spanning the secretory, nucleocytoplasmic, and endolysosomal trafficking pathways. This positions LRRK2 at the intersection of multiple essential pathways driving progerin pathology.

### LRRK2 reduction mitigates physiological aging in primary human cells and *C. elegans*

Given the shared molecular and cellular hallmarks between HGPS pathology and physiological aging, we hypothesized that *LRRK2* knockdown would similarly ameliorate aging-associated defects in cells from physiologically aged individuals ^8^. Knockdown of *LRRK2* in fibroblasts derived from multiple physiologically aged individuals (age range: 82-84 years) improved the aging status of multiple cellular aging markers, suggesting a broad pro-aging function of LRRK2 in physiological and progerin-induced cellular aging (Fig. 4A and B, S7A).

**Figure 4:**
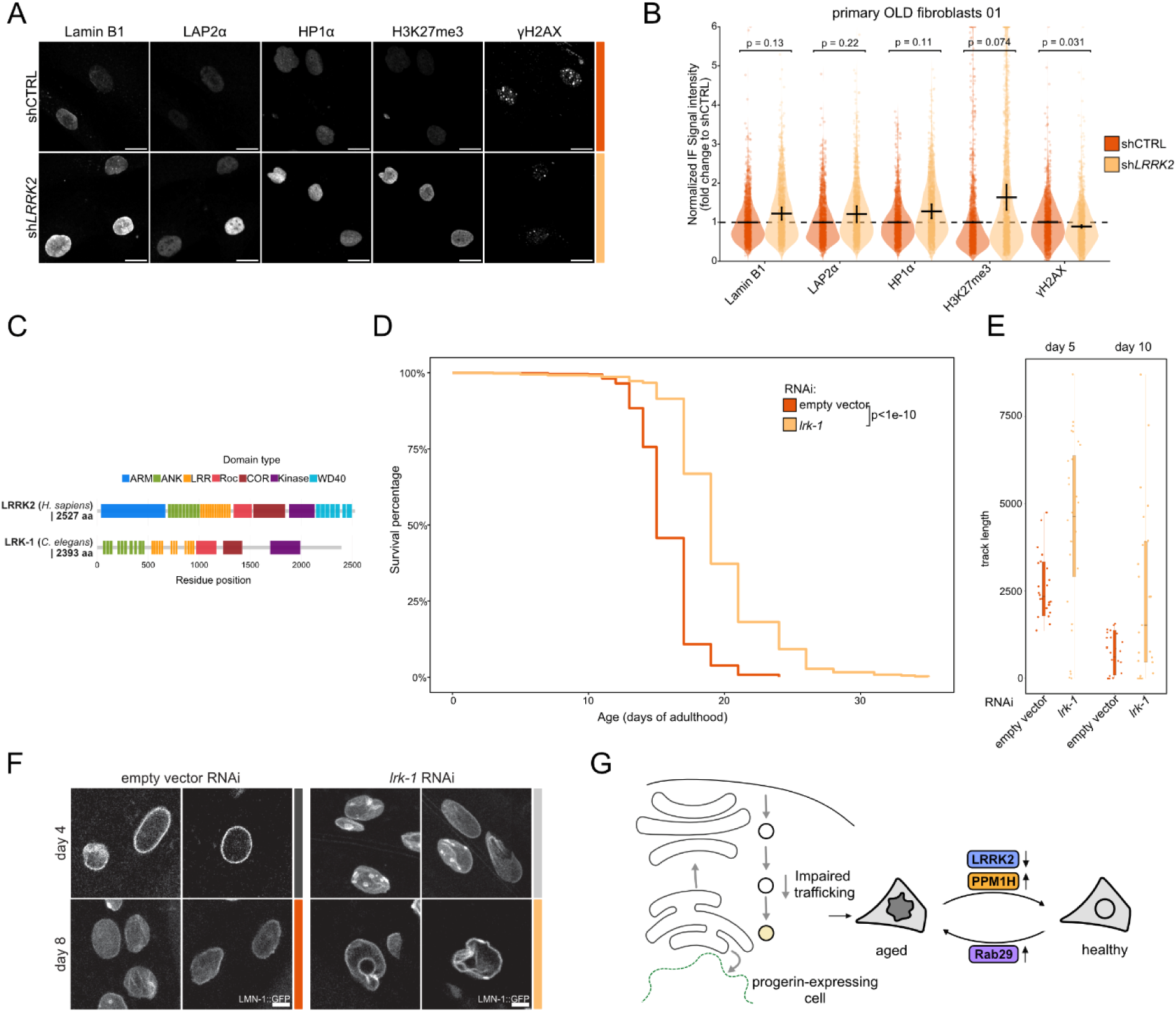
Knockdown of *LRRK2* and the *C. elegans* homolog *lrk-1* attenuates physiological aging defects. A, B: Representative immunofluorescence images (A) and quantification (B) of cellular aging markers in fibroblasts from physiologically aged donors expressing control or *LRRK2*-targeting shRNA. Scale bar: 20 μm. Values represent means ± SD (N=3). Statistical significance was assessed using a paired two-tailed Student’s t-test. C: Schematic of human LRRK2 and of *C.elegans* homolog LRK-1. Domains are depicted as follows: armadillo repeat (ARM), ankyrin repeat (ANK), leucine-rich repeat (LRR), Ras-of-Complex (ROC) GTPase, C-terminal-of-ROC (COR), Kinase, WD40 repeats ^26^. D: Representative survival curves of wild-type *C. elegans* treated with empty vector or *lrk-1* RNAi cultured at 20 °C. P-value (log-rank test) was calculated from N=3 with 90-120 worms per replica. E: Quantification of wild-type *C. elegans* movement treated with empty vector or *lrk-1* RNAi at the indicated days of adulthood. Movement was quantified as the length of tracks per minute (in pixels). N=3, p(day5)<2.2^-13, p(day10)<2.2^-13 (Wilcoxon-rank sum test). F: Representative single plane confocal images of hypoderm cells from *C. elegans* expressing *lmn-1::gfp* treated with empty vector, or *lrk-1* RNAi at days 4 and 8 of adulthood. Scale bar = 4 µm. G: Progerin expression induces defects in ER-to-Golgi protein trafficking, reduced endocytosis and lysosomal cargo transport, and altered import of nuclear proteins. Knockdown of the endolysosomal trafficking regulator LRRK2 or its counteracting phosphatase PPM1H ameliorates progerin-induced cellular aging, whereas LRRK2 activation via Rab29 promotes aging independently of progerin.

To test whether the beneficial effects of *LRRK2* knockdown we observed in cell culture models also translate to improvements on an organismal level, we performed RNAi knockdown of the *LRRK2* homolog *lrk-1* in *C. elegans*. *lrk-1* is highly evolutionarily conserved in its Roc-COR-kinase domain architecture and has a conserved function in *C. elegans* in regulating membrane trafficking (Fig. 4C) ^49^. Loss of *lrk-1* significantly extended the lifespan in *C. elegans* in comparison to worms treated with an empty vector control (Fig. 4D). To assess whether knockdown of *lrk-1* also improves health span, we quantified the aging-associated loss in mobility. While control worms exhibit a reduction in mobility between day 5 and day 10 of adulthood, *lrk-1* knockdown significantly rescued the decline, resulting in improved locomotor function in later life (Fig. 4E). Intriguingly, despite systemic benefits on life- and health-span, nuclei from *lrk-1* knockdown worms exhibit pronounced morphological alterations, including misshapen appearance and loss of sphericity (Fig. 4F, S7B and C). These findings indicate that *lrk-1* plays a role in maintaining nuclear envelope architecture, yet the anti-aging effects of *lrk-1* knockdown seem to outweigh any potential negative effects on the nuclear envelope. Collectively, we characterize defects in protein trafficking as part of HGPS cellular pathology and establish LRRK2/LRK-1 as a driver of HGPS pathology and physiological aging, making it a potential therapeutic target to ameliorate aging-associated defects (Fig. 4G).

## Discussion

Proteotoxic defects have emerged as central drivers of HGPS pathology. They are characterized by progerin aggregation, sequestration of key regulatory proteins, ER Stress, and impaired protein secretion ^11,16,19^. Here we describe widespread disruptions in cellular transport networks as part of HGPS pathology, including altered ER-to-Golgi trafficking dynamics, defects in endocytosis, and alterations in INM protein localization. Our findings identify the Parkinson’s disease-associated kinase LRRK2 as a promising therapeutic target for ameliorating HGPS pathology and reversing aging-associated defects, even in cells with established aging phenotypes.

Both knockdown of the *LRRK2* kinase and overexpression of its counteracting PPM1H phosphatase ameliorated progerin-induced cellular aging defects, while overexpression of the LRRK2 activator Rab29 or knockdown of *PPM1H* showed further deterioration of cellular aging markers. Our findings suggest that LRRK2 kinase activity drives pathological defects in protein trafficking, whereas reducing this activity or shifting the balance toward enhanced dephosphorylation of LRRK2 targets is beneficial in the context of cellular aging. The variable magnitude of rescue observed after *LRRK2* knockdown or PPM1H overexpression aligns with the inherent heterogeneity of cellular aging and HGPS and indicates that the beneficial effects depend critically on the extent to which the balance between LRRK2 and PPM1H is modulated. Nonetheless, our results show that reducing LRRK2 activity ameliorates cellular aging phenotypes in progerin-expressing cells.

Our findings on HGPS are interesting in the context of PD, where a regulatory imbalance of LRRK2 and PPM1H was previously reported to disrupt axonal transport of autophagosomes in neurons expressing PD-associated hyperactive LRRK2 mutants ^50^. We show that *PPM1H* knockout phenocopies the detrimental effects of hyperactive LRRK2, while PPM1H overexpression rescues defects of hyperactive LRRK2 mutants on axonal transport of autophagic vesicles ^50^. We suspect that the imbalance between LRRK2 and PPM1H is a common characteristic of HGPS and PD pathology. Mechanistically, this also indicates that the HGPS mechanism does not act through the kinase-independent Sec16A scaffolding function at Endoplasmic Reticulum Exit Sites (ERES) but rather through its canonical role in regulating Rab protein function ^51^. How precisely the numerous Rab proteins downstream of LRRK2 mediate this effect remains unclear. Unraveling this question is non-trivial, as the network of Rab proteins has multiple overlapping and, in some cases, counteracting functions ^52^. Nevertheless, our results indicate that Rab proteins downstream of LRRK2 are important regulators of progerin-induced cellular aging, further underlining that in the context of HGPS and cellular aging, LRRK2 kinase activity becomes a critical regulatory step and promising point of intervention. This is important as LRRK2’s focused therapeutic approaches are almost entirely kinase-activity–based (e.g., MLi-2-class inhibitors) ^53^.

We have no indication that LRRK2 levels or activity are altered by progerin, however, its role in protein trafficking connects it to most of the pathways that drive the loss of cellular homeostasis in aging. In support, in a PD mouse model, LRRK2 overexpression impairs Golgi apparatus organization, whereas its genetic ablation preserves Golgi structure ^43,54^. LRRK2 also colocalizes with ER and Golgi markers, and both its depletion and overexpression of hyperactive mutants disrupt ER-to-Golgi transport ^51^. Furthermore, LRRK2 phosphorylates critical regulators of vesicle budding and clathrin uncoating, while the expression of its hyperactive variants directly disrupts synaptic vesicle endocytosis in PD models ^55,56^. Additionally, PD-associated LRRK2 mutants cause lysosomal enlargement and reduced degradative capacities ^57,58^. At the same time, LRRK2 is required for maintaining mitochondrial function and can localize to the outer mitochondrial membrane ^59^. Cells expressing PD-associated LRRK2 mutants exhibit compromised ATP production, reduced mitochondrial activity, increased susceptibility to oxidative stress, and mtDNA damage ^60–62^. Beyond cytoplasmic organelles, LRRK2 directly interacts with lamin A/C and lamin B1 and B2 to maintain nuclear integrity ^63^. Disruption of this interaction, particularly by the expression of PD-associated mutants, leads to lamina disorganization, loss of nuclear integrity, and alterations in chromatin organization ^63,64^.

Aging is a multifactorial and tightly interwoven process, making it challenging to identify a single point of intervention that mitigates the detrimental effects associated with aging. Our data, together with previous reports, demonstrate that HGPS pathology is defined by similar defects observed in physiological aging and aging-associated diseases, such as impaired mitochondrial function, disrupted membrane trafficking and proteolytic pathways, and collapse of nuclear homeostasis ^6^. Given that LRRK2 centrally regulates these pathways, we propose that LRRK2 is a common denominator between physiological aging and progerin-induced aging. Since LRRK2 was already extensively investigated as a drug target in the context of PD with clinical trials currently in progress, we consider LRRK2 a very promising therapeutic target to ameliorate aging-associated defects in HGPS and physiological aging.

## Methods

### Cell culture and treatments

Human primary dermal fibroblast cell lines were obtained from The Progeria Research Foundation (PRF) Cell and Tissue Bank. The HGPS cell lines were HGADFN188, HGADFN178, HGADFN167; the control line was HGFDFN369. The following cell lines were obtained from the NIGMS Human Genetic Cell Repository at the Coriell Institute for Medical Research: AG11725, AG06292, AG14421, GM05659. In this study, the HGPS patient lines are designated as HGPS 01 (HGADFN188, donor age: 2 years and 3 months), HGPS 02 (HGADFN178, donor age: 6 years and 11 months), and HGPS 03 (HGADFN167, donor age: 8 years and 5 months). The cells from physiologically aged individuals are referred to as OLD 01 (AG11725, donor age: 84 years), OLD 02 (AG06292, donor age: 82 years), and OLD 03 (AG14421, donor age: 83 years). Primary fibroblasts were used after an additional 11–18 passages following their initial receipt. Additionally, apparently healthy control cell lines GM05659 (donor age: 1 year) and HGFDFN369 (donor age: 33 years and 9 months, only used in Fig. S2C) were hTERT immortalized, immortalized GM05659 cell line was used to generate doxycycline-responsive GFP-progerin-inducible fibroblasts. Previously generated progerin-inducible fibroblasts (P1 cells) were used for the siRNA screen (Fig. 1B) and in Fig. S1, S4C-E, S6B-C ^9^. HEK-293FT cells and CRISPRed HEK293T to Disrupt Antiviral Response (CHEDAR) cells were used to generate lentiviruses ^65^. All cell lines were grown in high glucose Dulbecco’s Modified Eagle Medium supplemented with 0.11 g/L Sodium Pyruvate (Gibco), 2 mM L-Glutamine (Gibco), 100 U/ml Penicillin-Streptomycin (Gibco), Non-Essential Amino Acids (Gibco) and either 10% (HEK293-FT/CHEDAR), 15% (P1) or 20% (primary and immortalized fibroblasts) sterile-filtered fetal bovine serum (FBS, Cytiva, SV30160.03) in a humidified incubator (Thermo Scientific) at 37 °C, 5% CO₂, and 90% relative humidity.

To induce GFP-progerin expression, GFP-progerin-inducible fibroblasts were incubated with 100 ng/ml doxycycline (dox, P1 cells 4 ug/ml doxycycline) for 96 hours, unless otherwise stated. For the Golgi inhibition experiment, P1 cells were cultured for 48 hours with vehicle (Ethanol) or GolgiStop diluted 1:1500 (BD Biosciences) in cell culture media. To release the fluorescent reporter in the RUSH assay, progerin-inducible fibroblasts expressing RUSH components were incubated with 100 ng/ml dox for 96 hours and treated 1 hour or 15 minutes before fixation with 40 μM Biotin (Sigma-Aldrich). For the Endocytosis assay, we induced P1 cells for 96 hours, serum-starved for 1 hour, treated for 30 minutes or 2 hours with 10 μg/ml Dil-LDL (Invitrogen), and washed 2 times with DMEM to reduce background before fixation.

### Cloning and generation of cell lines

Primary healthy fibroblasts were immortalized using pLenti CMV puro hTERT, or for experiments in Fig. 3B-E, where no additional antibiotic selection was possible with pLenti CMV hTERT constructs ^66^. rtTA3-based GFP-progerin-inducible cells were generated as previously described ^9^. In short, immortalized fibroblasts were sequentially transduced with lentivirus encoding pLenti CMV rtTA3 Hygro (Addgene, #26730) and pLenti CMV TRE3G Neo GFP-Progerin Dest (Addgene, #118710) ^9^. The blasticidin selection cassette from pLenti CMV TetR Blast (Addgene, #17492) was replaced with the Hygromycin cassette from pLenti CMV rtTA3 Hygro (Addgene, #26730) using *PfoI* and *NheI* enzymes (Thermo Scientific). To generate the pLenti CMV/TO Neo GFP-progerin construct, we transferred previously generated pENTR1A-GFP-Progerin into pLenti CMV/TO Neo DEST (Addgene, #17292) by LR recombination (Invitrogen) ^9^. We generated TetR-based GFP-progerin inducible cells by sequentially transducing immortalized fibroblasts with lentivirus encoding pLenti CMV TetR Hygro and pLenti CMV/TO Neo GFP-progerin. shRNAs targeting *LRRK2*, *PPM1H*, and a non-targeting control were cloned using *EcoRI* and *BamHI* sites (NEB) into a pSIH H1 Blast vector, a Blasticidin-resistant modified version of the pSIH H1 vector (System Biosciences). shRNA target sequences are listed in Supplementary Table 2. The selection cassette from the pCRISPRia-v2 (Addgene, #84832) plasmid was removed, and gRNA sequences were inserted using *BstXI* and *BlpI* sites (NEB). gRNA target sequences are listed in Supplementary Table 2. Cells were sequentially transduced with lentivirus encoding pLenti dCAS-VP64-Blast (Addgene, #61425) and modified gRNA-containing pCRISPRia-v2. FLAG-Rab29, FLAG-PPM1H, and V5-Rab (9A, 10, 12, 35, 43 (WT, CA, and DN mutants, exact mutations are listed in Supplementary Table 2) were cloned into a pMK vector containing attL sites. PPM1H construct was transferred into pLenti CMV Blast DEST (Addgene, #17451), and the RAB constructs into pLenti PGK Blast DEST (Addgene, #19065) by LR recombination. For the RUSH constructs, hooks and reporters were separately transferred into the pMK vector containing attL sites (Hook Str-STIM1-NN from Addgene, #65263, Hook Ii-Str from Addgene, #65289, Reporter SBP-mCherry-GPI from Addgene, #65295, Reporter TNFα-SBP-mCherry from Addgene, #65279). Reporter and Hooks were transferred into pLenti CMV Puro Dest (Addgene, #17452) and pLenti CMV Blast Dest (Addgene, #17451), respectively, by LR recombination. Cells were transduced with lentivirus as previously described using HEK-293FT cells or CHEDAR cells when higher virus titers were required ^67^. 48 hours post-infection, cells were selected using Puromycin (Gibco), 37 ug/ml Hygromycin (Gibco), 333 ug/ml Neomycin (Gibco), or 6.1 ug/ml Blasticidin (Gibco).

### Immunofluorescent Staining and Imaging

All steps for IF staining were performed at room temperature. Washing steps were carried out using a Blue Washer (BlueCatBio). Cells were seeded and induced as appropriate in 384-well plates (Revvity). 96 hours post-induction/seeding, cells were fixed using 4% formaldehyde/PBS (Thermo Scientific) for 10 minutes, followed by one wash with PBS/0.05% Tween-20 (Sigma-Aldrich). Next, fixed cells were permeabilized for 10 minutes (PBS/0.5% Triton-X 100 (Sigma-Aldrich)), followed by one wash with PBS/0.05% Tween-20. Primary antibodies were diluted in blocking buffer (PBS, 0.05% Tween-20, 5% bovine serum albumin (BSA, Sigma-Aldrich)), and cells were incubated for 1 hour with primary antibodies. Afterwards, cells were washed twice with PBS/0.5% Tween-20 and once with PBS/0.05% Tween-20. After the washes, cells were incubated for 1 hour with secondary antibodies with 2 μg/ml DAPI and washed again twice with PBS/0.5% Tween-20 and once with PBS/0.05% Tween-20. Cells were kept at 4 °C until imaging. For endocytosis tracking, cells were fixed similarly, washed once with All-purpose solution (APS, 0.03% Saponin (Sigma-Aldrich), 0,2% BSA in PBS), and incubated for 30 minutes in APS to block. Afterwards, cells were incubated for 1 hour with primary antibody diluted in APS, washed 3 times with APS, incubated with secondary antibodies diluted in APS for 1 hour, washed 3 times again with APS, and stored at 4 °C in PBS until imaging. Primary and secondary antibodies used for immunofluorescence are detailed in Supplementary Table 2. Images were acquired using an Opera Phenix High-Content Screening System (Revvity) in confocal mode with a 40x/1.1 NA water-immersion objective or 20x /1.0 NA water-immersion objective (RUSH assay) in three sequential steps (405/640 nm, 488 nm, and 561 nm excitation lasers, emission filters: 405 nm excitation: 435 nm - 480 nm; 640 nm excitation: 650 nm -760 nm; 488 nm excitation: 500 nm -550 nm; 561 nm excitation: 570 nm to 630 nm). Typically, 12-16 single-plane images were acquired per well, 4 wells per condition and per antibody were imaged. For the RUSH assay, 9 single-plane images were acquired per well, 4 wells per condition and per antibody were imaged. For the quantification of ER and Golgi morphological features, cells were imaged as Z-Stacks and analyzed as 3D objects. Images were analyzed and quantified using Columbus 2.9.1 (Revvity) and Harmony 5.2 (Revvity) and prepared for publication using Fiji ^68^. Nuclei were segmented based on DAPI staining, cytosol was segmented using cytosolic vimentin staining, and Golgi and ER segmentation were based on GRASP65 and BIP staining. For the detection of endocytic compartments, we used EEA1 (Early Endosomes), Rab7 (Late Endosomes), and LAMP2 (Lysosomes) staining for segmentation. We quantified the mean nuclear intensities of GFP-progerin, lamin B1, LAP2α, HP1α, H3K27me3, and H3K9me3, and foci intensity of γH2AX and 53BP1. For the RUSH-assay, we quantified the accumulation of the fluorescent reporters within the segmented Golgi apparatus. Similarly, we quantified Dil-LDL intensity in positive spots for early endosome, late endosome, and lysosomal markers for endocytosis tracking. In the plots, individual cells are depicted as fold change relative to the mean of the control/uninduced control condition. To quantify the cytosolic localization of nuclear envelope proteins, a cytoplasmic ring region surrounding the nuclei was generated from the nuclear mask, and the mean fluorescence intensity within this ring was measured. Nuclear roundness was calculated in Harmony as sqrt(4*pi*Area)/Perimeter, with 1 representing a perfect circle and lower values indicating reduced nuclear circularity. The complete Harmony analysis pipelines can be found in Supplementary Table 3.

### RNA isolation and analysis of gene expression

Total RNA was extracted using TRIzol (Invitrogen) according to the manufacturer’s instructions. Briefly, cells were lysed in 500 µL TRIzol, followed by phase separation with chloroform (Roth) and RNA precipitation with isopropanol (Honeywell). The RNA pellet was washed with 70% ethanol (Roth), air-dried, and resuspended in DEPC-treated water. The RNA was transcribed into cDNA by reverse transcriptase (In-house) using random hexamer primers (Thermo Scientific), and qPCR was performed using Power SYBR Green PCR Master Mix (Thermo Scientific), using 40 amplification cycles (95 °C 10 seconds, 62 °C 15 seconds, 72 °C 10 seconds) and a Roche LightCycler 480II. We performed all qPCRs using *GAPDH* as a housekeeping gene to normalize gene expression. Primer sequences are listed in Supplementary Table 2.

### Western Blot

Cells were harvested using 0.05% Trypsin-EDTA (Gibco) and snap-frozen on dry ice. Next, cell pellets were resuspended in sample loading buffer (1x Laemmli Sample Buffer (Bio-Rad), 5% 2-Mercaptoethanol (Sigma-Aldrich), 1x PBS) and boiled at 95 °C for 5 minutes. Extracts were loaded on Criterion TGX Precast 4-15% gradient gels (Bio-Rad), transferred to a 0.2 μm PVDF membrane (Bio-Rad), and blocked with 5% skimmed milk (Millipore, in PBS-T (0.1% Tween-20)) for 1 hour. The blocked membrane was incubated with primary antibodies diluted (see Supplementary table for details) in 5% milk (primary mouse antibody) or 5% BSA (primary rabbit antibody) overnight at 4 °C. Afterwards, membranes were washed 3 times for 5 minutes in PBS-T and incubated for 1 hour with HRP-labeled secondary antibody diluted in milk. After 3 more washes for 5 minutes in PBS-T, blots were kept in PBS until development. For the detection of phosphorylated proteins, PBS was replaced with TBS. Immunoluminescence of the secondary antibodies was detected with ECL detection reagent (Thermo Scientific) on a Bio-Rad ChemiDoc XRS+. Blots were analyzed using the Image Lab (Bio-Rad) and quantified using Fiji ^68^. Antibodies used for Western blotting are detailed in Supplementary Table 2.

### Kinome-wide siRNA Screen

The kinome-wide siRNA screen was performed with P1 cells using an NCATS in-house kinome library targeting 704 human genes, each with 3 individual siRNAs, and was done in dual replicates. Cell plating, staining, imaging, and analysis were done as previously described ^11^. Hits were identified based on MAD fold change (MAD FC) to control siRNA in the progerin ON condition. siRNAs were considered at hit if MAD FC GFP-progerin < -2.5, MAD FC lamin B1 > 2.75, MAD FC γH2AX < -2. siRNAs with an MAD FC < -1.5 for cell numbers were excluded.

### *C. elegans* maintenance, RNAi feeding, and lifespan assays

Wild-type *C. elegans* (N2 (Bristol)) was grown at 20 °C and synchronized by bleaching gravid adults. L1 was grown by hatching embryos at room temperature in M9 buffer. The synchronized L1s were transferred and grown until the L4 larval stage in NGM plates containing 4 mM IPTG and 100 μg/ml carbenicillin (RNAi plates) and seeded either with HT115 bacteria expressing the empty vector (L4440) or dsRNA against *lrk-1* (both obtained from the Ahringer library and verified by sequencing). For each condition, ∼100 late L4 larvae were transferred to 12-well plates with 9-15 worms per plate. Each well contained 1 ml of NGM supplemented with Carbencillin (100 μg/ml), IPTG (4 mM), and 5-Fluoro-2′deoxyuridine (50 μM) (FudR, Sigma-Aldrich) and was seeded with the respective RNAi-expressing bacteria. The L4 stage was designated as day 0 of the analysis. Worm lifespan was assessed by counting moving worms every 2-3 days using a stereo dissection microscope (Leica M80). Animals that crawled off the plate were excluded from the analysis. The data were plotted with Kaplan-Meier Survival curves. For pairwise comparison, the log-rank Mantel–Cox statistical test was used. Around 50 synchronized wild-type (N2 Bristol) larvae were transferred to 4 plates per replica and RNAi condition, seeded with HT115 bacteria expressing *lrk-1*, or empty vector (L4440) dsRNA. Brightfield time-lapse movies (1 minute total duration, 2 seconds interval) were recorded using a Zeiss AXIO Zoom v16 at the indicated time points. To ensure movement, brightfield movies were preceded by 5 seconds interval of continuous blue light. To quantify speed of movement, we segmented and tracked individual *C. elegans* using machine learning algorithms implemented in Ilastik (https://www.ilastik.org/), the pixel classification module followed by the Ilastik animal tracking workflow. Final speed was plotted as track length per minute, in pixels. Around 50 synchronized larvae expressing *lmn-1::gfp* were transferred to RNAi plates seeded with HT115 bacteria expressing *lrk-1*, or empty vector (L4440) dsRNA ^69^. On days 4 and 8 of adulthood, worms were mounted on agarose pads with (−)-Levamisole hydrochloride solution (1 mM) (Sigma-Aldrich) and imaged with the Stellaris 8 FALCON confocal microscope (Leica).

